# Endogenous circannual cycles drive seasonal cold hardening in temperate ants

**DOI:** 10.1101/2025.07.09.663877

**Authors:** Quentin Willot, Vladimír Koštál, Johannes Overgaard

## Abstract

The ability to acquire cold tolerance and cope with seasonality underlies the broad geographical distribution of ectotherms. Among insects, cold-adapted ants, which are successful terrestrial insects in temperate and boreal ecosystems, rely on complex circannual life cycles to survive multiple consecutive winters. Following winter reactivation, colonies enter a fixed developmental phase whose duration is controlled by “Kipyatkov’s sand-glass device”, an endogenous programmed timer that ultimately enforces a new dormancy period for colonies after a set period, even under permissive light or temperature cues. Here, using five temperate ant species, we leveraged this system to tease apart the contribution of intrinsic seasonal programming and extrinsic thermal cues on worker cold tolerance, metabolic rates, and metabolomic profiles. We show that obligate colony-level dormancy alone is sufficient to modulate both the critical thermal minimum (CT_min_) and the temperature causing 50% mortality during acute cold exposure (LTe_50_), and further interacts with cold acclimation to shape worker phenotypic cold tolerance. While colony-level dormancy was only associated with limited metabolomic reorganization, cold acclimation triggered the accumulation of metabolites potentially involved in osmotic balance and membrane reorganization. Our results highlight a circannual cycle of endogenous cold hardening that operates independently of environmental exposure in temperate ants, providing new insight into the physiological adaptations underpinning the evolutionary success of this insect family in highly seasonal climates.

## Introduction

Seasonal temperate climates pose several challenges to ectotherms. Winter is typically associated with reduced energy availability and exposure to potentially lethal cold temperatures^1^. In insects facing such challenges, these environmental constraints have led to the emergence of complex metabolic dormancy phenotypes such as diapause and quiescence^2–5^. Diapause is a stage-specific and programmed developmental arrest characterized by metabolic suppression^2–7^. It is often triggered by environmental cues in advance of the actual challenging conditions (facultative diapause, e.g., signaled through shortening days), and in some species, can also be genetically encoded to occur at a specific life stage regardless of environmental conditions (obligate diapause)^2–5^. In contrast to diapause, quiescence is a form of dormancy that occurs only as an immediate response to limiting environmental factors (low temperature, dehydration, fasting, hypoxia, etc.)^3–5^. Numerous studies have associated both winter diapause and quiescence with energy saving and increased resilience towards environmental stress in cold-adapted invertebrates^3,5–9^. However, the direct relationship between acquisition of cold tolerance and programmed dormancy in insects remains poorly understood^3,10^ In some species, diapause triggers the preemptive buildup of cryoprotective molecules^11–15^, indicating at least partially overlapping pathways^3^. In others, despite diapause, accumulation of cryoprotective molecules occurs only after exposure to low temperatures, implying cold hardening and diapause to be mainly coincidental^16,17^.

Ants offer an opportune model system to study the interactions of seasonal biological cycles and cold tolerance. Ancestrally tropical, ants have successfully radiated and adapted to most seasonal climates on the planet^18,19^. Further, individuals are usually long-lived^20^, thus they have to endure several consecutive winters over their lifespan. For example, queens of the European black garden ants *Lasius niger* can reach up to 28 years in captivity and workers of the same species display an estimated lifespan of roughly 1 to 3 years^21^. At the colony-level, many temperate species are characterized by annual growth cycles that involve developmental arrest at collective level (colony dormancy), potentially coinciding with individuals’ diapause. As described by Kipyatkov and colleagues, three main broad categories of annual colony-level growth cycles have been identified in ants (Fig. S1)^22–25^. First, tropical ant species appear unable to slow or halt colony-level growth in response to external cues (homodynamic colony cycle); workers are chill-sensitive and only capable of limited quiescence^22–25^. In contrast, subtropical and temperate species are able to enter either facultative (exogenous-heterodynamic) or obligate (endogenous-heterodynamic) prolonged periods of colony-level dormancy linked to higher winter survival (Fig. S1)^22^. Furthermore, in species displaying obligate diapause (endogenous-heterodynamic), the duration of the yearly developmental phase of colonies (from dormancy to dormancy) appears largely governed by an endogenous timer with only limited sensitivity to external conditions (such as temperature and/or day length)^22,23,26–28^. Captive colonies invariably enter obligate dormancy after a finite time interval, with environmental factors able to delay but not prevent its onset (Fig. 1A)^22^. This phenomenon led Kipyatkov to propose the existence of a “Sand-glass device” in these species, a master switch governing the length of seasonal colonial development based on endogenous colony-cycles, with minimal integration of environmental cues (Fig. 1A)^22–25^. Despite their success in temperate ecosystems, how colony-level dormancy and environmental exposure integrate to modulate the cold tolerance of temperate ants remains unexplored.

**Figure 1.**
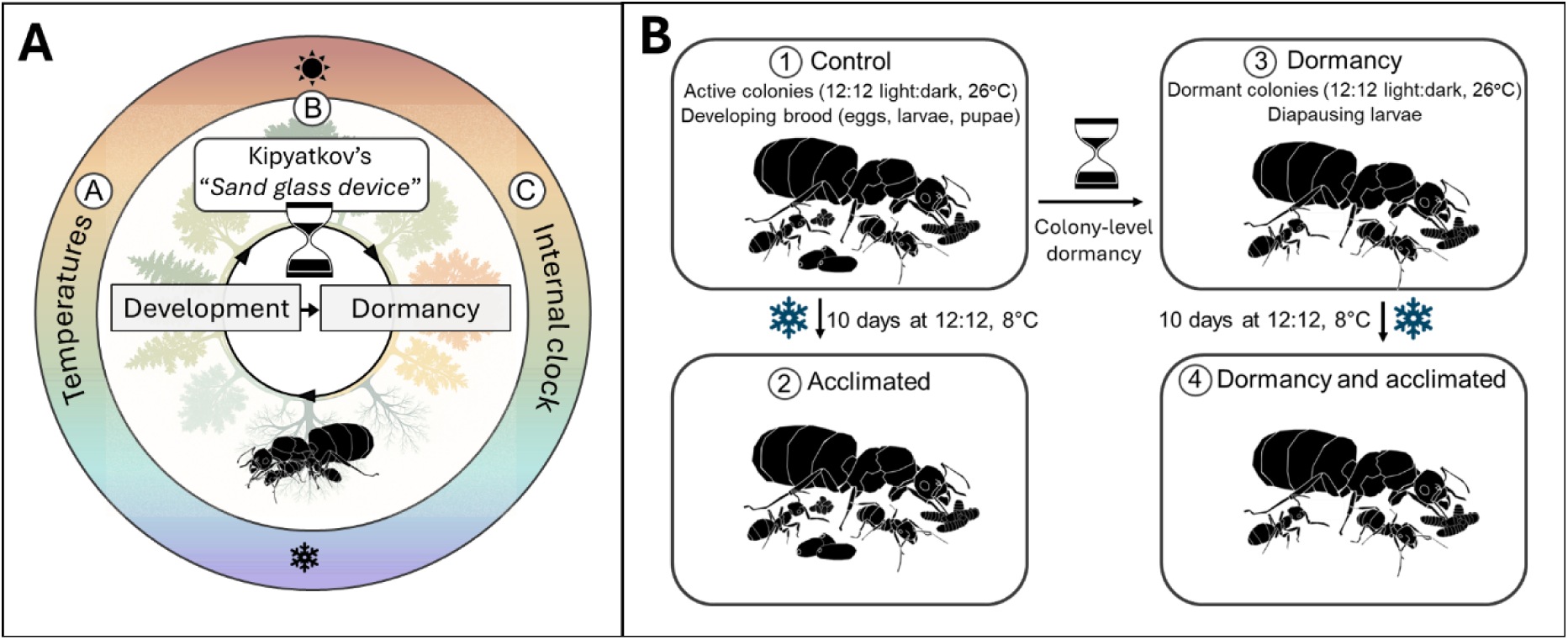
**A.** Yearly developmental cycle of endogenous-heterodynamic ants. A. Environmental cues, such as rising temperatures, trigger the end of winter dormancy for colonies in spring. B. Colonies then enter a fixed growth period, with queen egg-laying, worker activity, and brood development. C. Regulated by Kipyatkov’s “*Sand Glass Device*”, an internal clock determines the onset of the next dormancy with only limited integration of environmental cues (e.g. dormancy is initiated after a fixed period even when ants are exposed to warm temperatures or long days), typically coinciding with late summer or early fall locally. This phase is marked by the cessation of queen egg-laying, reduced worker activity, and depending on the species, the presence of diapausing larvae whose development will resume next spring. **B.** Experimental design. We leveraged the developmental cycle of endogenous-heterodynamic ants to establish a full-factorial design disentangling the relative effects of (i) acclimation and (ii) colony-level dormancy on the cold tolerance and metabolic reorganization of workers.

Here, we disentangle the relative effects of colony-level dormancy and acclimation on worker cold tolerance. We used five common endogenous-heterodynamic European ant species spread across four distantly related genera, and leveraged a full-factorial design based on dormant vs non-dormant colonies and cold acclimated vs non-cold acclimated individuals. Through this system, we measured the (i) lower limits for neuromuscular activity (Critical Thermal Minimum, CT_min_), (ii) acute limits of cold-stress mortality (LTe_50_), (iii) metabolic rates putatively indicative of diapause, and finally (iv) the metabolomic reorganization hypothetically associated with increased cold tolerance in workers.

## Methods

### Species sampling, husbandry and acclimation

Colonies of *Lasius niger*, *Lasius flavus*, *Formica fusca*, *Myrmica rubra,* and *Leptothorax acervorum* were collected near Aarhus, Denmark (56.16°N, 10.20°E) in summer 2021. These four genera encompass the majority of species diversity in subboreal and boreal ecosystems, accounting for around 80% of the native specific ant diversity found in Denmark^29^. Colonies were brought back to a climatized chamber and kept in plastic boxes with Fluon®-coated sides and provided with 16×150 mm plastic test tubes with a water-filled section in the bottom separated with a moist cotton plug for nesting. During the developmental phase of the colony, characterized by worker foraging activity, queen egg-laying, and developing brood^22^, colonies were kept under a 12:12 light:dark cycle at constant 26°C, and fed honey water and sliced mealworms twice a week. At the onset of colonies’ dormancy, characterized by the absence of worker foraging activity, of queen egg-laying, and diapausing larvae with halted development (*Lasius*, *Myrmica*, and *Leptothorax* species) or no diapausing larvae at all (*Formica fusca*) ^22^, colonies were moved into a cold room kept under a constant 12:12 light:dark cycle at 8°C for 4 months. Following this cycle, all colonies were simultaneously returned to developmental conditions (constant 12:12, light dark cycle at 26°C) where queen egg-laying, foraging activity and larval development usually resumed in colonies within 1 to 2 weeks. Experiments began in 2022 following colonies’ first artificial laboratory dormancy cycle. As a result, data are reported for workers exposed solely to the conditions defined in the husbandry methods for a full year (dormancy to dormancy).

### Experimental design

We leveraged the endogenous-heterodynamic annual developmental cycle of the selected ant species to establish four experimental conditions (Fig. 1B). First, (1) control conditions, corresponding to workers taken from colonies in their active developmental phase, characterized by queen egg-laying and brood growth, maintained at a constant 12:12 light:dark cycle and 26°C. Second (2), acclimated conditions, corresponding to workers originating from the same control conditions but subsequently acclimated to 8°C for 10 days. Third, (3) dormancy conditions at warm temperature, corresponding to workers from the same colonies as control conditions but sampled later in their annual developmental cycle after the onset of colony-level dormancy (characterized by the absence of worker foraging activity, of queen egg-laying, and diapausing larvae with halted development or no diapausing larvae for *Formica fusca*^22^). Finally (4), cold-acclimated and dormancy conditions, corresponding to workers from dormancy conditions that had been moved to 8°C for 10 days. These four conditions allowed us to exploit a full factorial design disentangling the relative effects of (i) acclimation and (ii) colony-level dormancy on the cold tolerance and metabolic reorganization of workers (Fig. 1B).

### Dynamic cold tolerance assays

To investigate the lowest temperature limits for worker neuromuscular activity, Critical thermal minimum (CT_min_) of workers were recorded as described previously^30^. Briefly, workers were placed individually into 5 mL closed glass vials mounted to a rack and submerged into a transparent tank filled with a 25% ethylene-glycol:water solution. Temperature in the bath was initially maintained at 20°C for 15 min before it was gradually decreased at a rate of 0.1°C/min using a programmable water bath (LAUDA-Brinkmann, NJ, USA). Chill-coma (CT_min_) was then scored for each individual (N=10-20 workers per species) as the temperature resulting in a total absence of movement of workers (even after gentle vial shaking).

### Lower Lethal temperature

To investigate mortality to acute cold stress in workers, Lower Lethal temperatures (LTe_50_), defined as the temperature where 50% of individuals died following an acute cold exposure of 24h, were measured as previously described for other insects^31^. For each species and each condition, 35-mL vials containing 10-20 workers were placed for 24 h at a range of constant stressful temperatures. Each species was exposed to 6–12 different temperatures to obtain mortality data spanning from 100% to 0% (between all treatments and species this included temperatures ranging from +8 to −20 °C). After the 24h cold exposure, vials were returned to room temperature (ca. 21°C), and workers that were not able to move after an additional recovery period of 24h were scored dead. The temperature corresponding to 50% mortality (LTe_50_) was calculated by fitting a sigmoidal curve to the mortality data for each species and condition (see data analysis).

### Metabolic rates measurements

To investigate the relationship between colony-level dormancy and suppressed metabolic rates in workers, indicative of diapause, standard metabolic rates (SMRs) of our 5 species of ants were recorded across two conditions: (i) control, (workers taken from colonies in their developmental phase) and (ii) dormant (workers from the same colonies but sampled after the onset of colony-level dormancy). SMRs were estimated indirectly from the rate of CO_2_ production (VCO_2_) using repeated measurements of stop-flow respirometry as described previously for ants^30^. Measurements were made at 18°C as a conservative tradeoff between detectable metabolic signal and lower level of worker activity. On completion of measurements, the total fresh weight of workers was measured to the nearest 0.1 mg (Sartorius, Type 1712, Göttingen, Germany) to allow for calculation of SMRs.

### Metabolomics

To link modulation of cold tolerance across our experimental design with putative accumulation of cryoprotective molecules in workers, we sampled ants corresponding to our 4 experimental conditions (see design) to analyze their metabolomic profiles. Workers were snap-frozen in liquid nitrogen and their gaster (the portion of the abdomen containing the crop) was removed to avoid potential contamination from food content. We sampled from 5 species x 4 treatments x 4 replicates, except for *L. acervorum* where too little biological material was available to analyze control conditions. This resulted in 76 analytical samples. Samples were transported on dry ice to Biology Center, České Budějovice for analysis, and the fresh mass of each analytical sample was measured to the nearest 0.1 mg using a balance MSE6.6S (Sartorius), resulting in 5 – 10 mg of worker tissues pooled for analysis for each analytical sample. To prepare analytical samples, they were melted on ice and homogenized in 500 µL of extraction buffer – methanol:acetonitrile:deionized water (2:2:1, v/v/v; Fisher Scientific, Pardubice, Czech Republic). Twelve additional blanks without ant tissues (4 per experimental condition) were also prepared for subsequent analysis of background noise to the signal. Internal standards, p-fluoro-DL-phenylalanine, and methyl-D-glucopyranoside (from Sigma-Aldrich, Saint Louis, MO, USA) were added to the extraction buffer to standardize measurements, both at a final concentration of 200 nmol mL^−1^. Downstream metabolite extractions and analysis of 82 targeted metabolites were then performed as previously described^32^. Of the 82 initially targeted metabolites, 19 were not detected in our samples, leaving 63 metabolites for downstream analysis. A manual quality control (QC) check was then performed to curate the dataset: first, potential blank contamination values were subtracted from the analytical samples. Second, metabolites with a significant standard deviation (>33%) across replicates for any condition in one or more species were excluded. After these steps, 53 differentially expressed metabolites passed the QC check. These metabolites were quantified from their areas under their respective chromatographic peaks and then normalized to the amount of fresh mass of the respective analytical samples.

### Data analysis

Data analysis was performed in R version 4.1.2^33^. The comparison of CT_min_ values between our four experimental conditions for each species was performed using one-way ANOVAs followed by Tukey’s multiple comparison post-hoc tests. Assumptions of normality and homogeneity of variance were verified using the Shapiro–Wilk test and Bartlett’s test, respectively. LTe₅₀ values were computed by fitting sigmoidal curves to mortality proportions for all conditions and species, providing the temperature of 50% mortality after a 24h exposure (LTe₅₀). LTe₅₀ values are reported in the manuscript with their 95% confidence intervals (CIs), non-overlapping 95% CIs were considered indicative of statistically significant differences between conditions. As a redundant analytical step to assess the effects of (i) colony-level dormancy and (ii) acclimation on both CT_min_ and LTe₅₀ across our entire dataset, a model-building approach was implemented using the lmer() function from the lme4 package^34^ (after verifying the assumptions of a linear mixed-effects model). Model testing through Akaike Information Criterion (AIC) was performed using the aicw function from the package geiger^35^. The effect size of fixed variables included in the best-performing models (dormancy and acclimation) on CT_min_ and LTe₅₀ values was calculated through a partial Omega Squared statistic (ωp^2^) using the effsize package^36^. To test whether the onset of colony-level dormancy was associated with suppressed metabolism indicative of diapause in workers, we used a two-way ANOVA testing for the impact of species, dormancy, and their interaction on worker SMR, followed by a Tukey post-hoc test to assess pairwise differences between control and dormant conditions within each species.

To provide a holistic examination of dormancy status and acclimation on the metabolomic profiles of workers, we first performed a visual investigation of metabolomic data based on heat-map profiles of area under the chromatographic peak of metabolites’ fold change (FC) across conditions. Second, we performed a Principal Component Analysis (PCA) of metabolites differentially expressed across conditions using the function prcomp from the R stats package^33^. This PCA investigated within a single analysis which relative fluctuation of metabolites (area under the chromatographic peak) were generally associated with fluctuations in cold tolerance (CT_min_ and LTe_50_) across all species and conditions within our dataset.

## Results

### Endogenous onset of colony dormancy influences the CT_min_ of workers even in the absence of cold acclimation

We recorded CT_min_ values of workers across our 4 conditions to disentangle the relative effect of colony-level dormancy and cold acclimation on the cold tolerance of individuals. Workers from control conditions exhibited the highest CT_min_, with values ranging from 1.7°C (*L. niger*) to 0.2°C (*M. rubra* – Fig. 2 A-E). A cold acclimation period of 10 days at 8°C of workers from control, non-dormant conditions significantly lowered their CT_min_ across all species, indicating that CT_min_ values were reduced with cold acclimation. The onset of colony-level dormancy also lowered CT_min_ values across all species (Fig. 2), indicating phenotypic changes linked to cold tolerance in workers even in the absence of cold acclimation. Across the five species we found that the isolated effects of dormancy and acclimation were synergistic such that the lowest CT_min_ values were observed for workers having been cold-acclimated at 8°C for 10 days after the onset of colony-level dormancy, with CT_min_ values ranging from −3.2°C (*L. flavus*) to −4.3°C (*L. acervorum*) (Fig. 2A–E). The best fit model testing for the impact of both dormancy and acclimation on CT_min_ values identified through AIC comparison (Table S1) included both acclimation and dormancy, as well as their interaction, as factors (Table 1). Thus, the mixed-effect regression model confirmed that acclimation and colony-level dormancy were significant predictors of CT_min_ across species, independently of each other (p < 0.001; Table 1) but also with a strong interactive effect (p < 0.01; Table 1).

**Figure 2.**
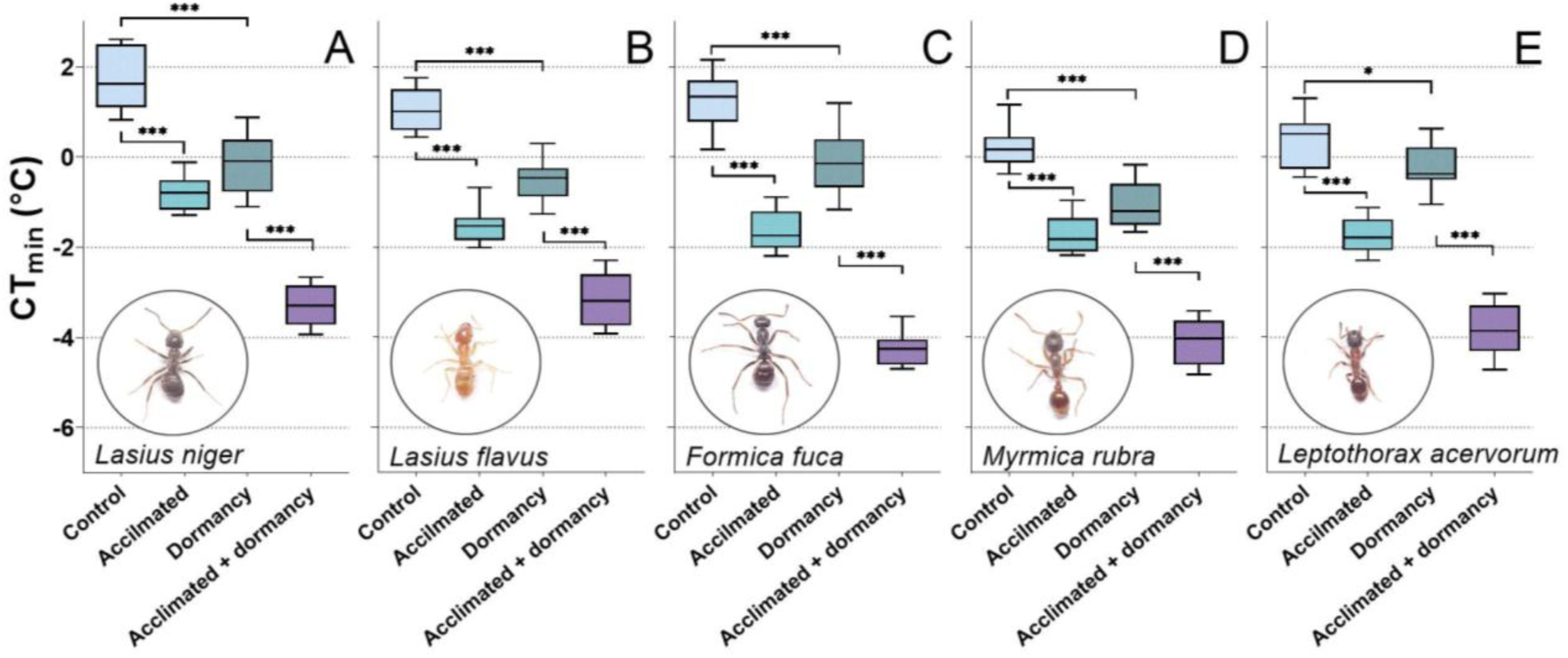
Effects of cold acclimation and endogenous colony-level dormancy on CT_min_ values of individuals across studied species. CT_min_ values (mean ± SD) were calculated for 10–20 individuals per condition and species. Both a cold-acclimation period and the onset of colony-level dormancy independently exerted a significant effect on CT_min_ values as compared to workers from control conditions (growing colonies maintained at 26°C, 12:12 light/dark cycles), while further displaying a strong and significant interactive effect (Table 1). Statistical significance between treatments within species is indicated (p < 0.05*, *p*< 0.01**, *p*< 0.001***; one-way ANOVA and Tukey *post hoc* test).

**Table 1.** Significance (*p*) and effect size (partial omega-squared values, ωp^2^) of each variable included in the best fit linear mixed-effects regression models teasing apart the effect of acclimation and colony-level dormancy on CT_min_ and LTe_50_ values (See table S1 for best model selection). Overall, both acclimation and dormancy statistically affected CT_min_ across species, yielding moderate to large effect sizes. In addition, these fixed effects displayed notable interaction, indicating that the effect of acclimation on CT_min_ was more pronounced for species after the onset of colony-level dormancy. In contrast, LTe_50_ values were significantly influenced across all species by acclimation only, with moderate effect size.

| CT<sub>min</sub> model | R<sup>2</sup> | Fixed Variables | *p* | Effect size<br>(ωp<sup>2</sup>) | Effect size 90%<br>one-sided CI |
| --- | --- | --- | --- | --- | --- |
| CT<sub>min</sub> ~ acclimation * dormancy + (1 \| sp) | 0.96 | Acclimation | 1.43*10<sup>-7***</sup> | 0.89 | 0.80-1 |
|  |  | Dormancy | 1.97*10<sup>-6***</sup> | 0.84 | 0.70-1 |
|  |  | Acclimation:dormancy | 6.29*10<sup>-3**</sup> | 0.41 | 0.14-1 |

**Table 1.** Significance (*p*) and effect size (partial omega-squared values, ωp<sup>2</sup>) of each variable included in the best fit linear mixed-effects regression models teasing apart the effect of acclimation and colony-level dormancy on CT<sub>min</sub> and LTe<sub>50</sub> values (See table S1 for best model selection).
| LTe<sub>50</sub> model | R<sup>2</sup> | Fixed Variables | *p* | Effect size<br>(ωp<sup>2</sup>) | Effect size 90%<br>one-sided CI |
| --- | --- | --- | --- | --- | --- |
| LTe<sub>50</sub> ~ acclimation + dormancy + (1 \| sp) | 0.71 | Acclimation | 1.91*10<sup>-4***</sup> | 0.67 | 0.45-1 |
|  |  | Dormancy | 0.07 | 0.23 | 0.02-1 |

### Tolerance to acute cold stress is primarily driven by cold acclimation

The proportion of dead ants following acute 24-hour cold stress was measured as a function of temperature across our five species for each of the four conditions (Fig. 3). The temperature causing 50% worker mortality over 24 hours was extracted from the fitted sigmoidal relationships to provide LTe_50_ values (Fig. 3, mean ± 95% CI). Workers from control conditions (26°C with a 12:12 light/dark cycle) exhibited the highest LTe_50_ values and were thus the most sensitive to acute cold stress, displaying LTe_50_ values ranging from −2.2°C (*L. niger*) to −9.5°C (*L. acervorum*). A 10-day cold acclimation at 8°C of workers from control conditions significantly lowered their LTe_50_ across all species (Fig. 3). The onset of colony-level dormancy also impacted LTe_50_ values when investigating species-specific patterns (Fig. 3A–E), although this effect was less pronounced than acclimation and not significant for *L. flavus* (Fig. 3B). There was limited support for additive/synergistic effects of dormancy and acclimation on LTe_50_ values when comparing dormant and non-dormant individuals, except for *F. fusca* (Fig. 3C). Overall, the lowest LTe_50_ values and lowest mortality proportions to acute cold stress were observed in acclimated or acclimated + dormancy workers, with values ranging from −8.5°C (*M. rubra*) to −13.5°C (*L. acervorum*). The best fit model testing for the impact of both dormancy and acclimation on LTe_50_ values identified through AIC comparison (Table S1) included both acclimation (p < 0.001) and dormancy (p = 0.07) as factors, without their interactive effect, and with dormancy approaching statistical significance only (Table 1).

**Figure 3.**
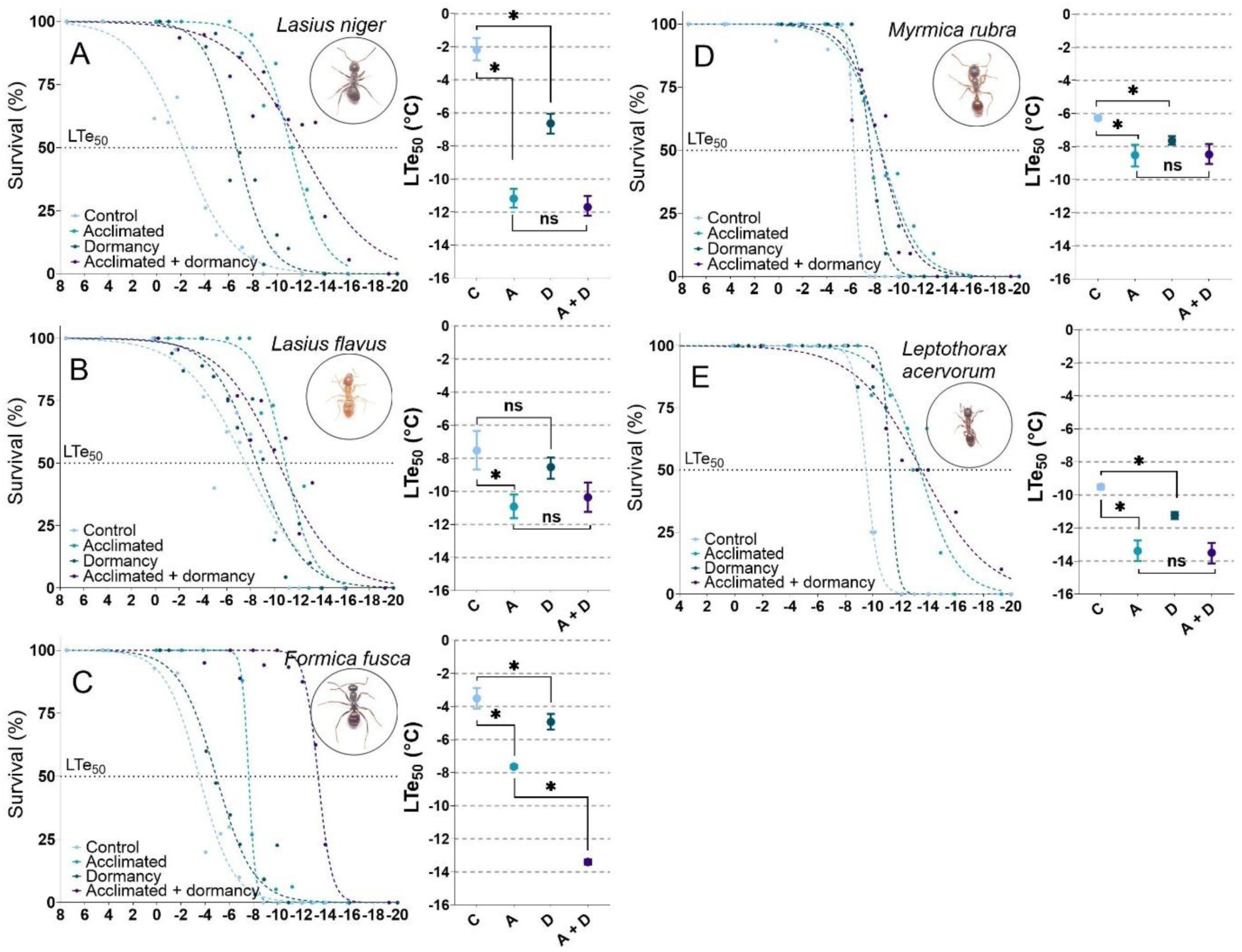
Effects of cold acclimation and colony-level dormancy on survival to acute cold-stress (LTe_50,_ mean ± 95% CI). **A.** *Lasius niger*. **B.** *Lasius flavus*. **C.** *Formica fusca*. **D.** *Myrmica rubra*. **E.** *Leptothorax acervorum*. For each species, 10-20 workers were placed at a single temperature ranging from +8 to −20 °C for 24h, and survival was scored after an additional 24h recovery period to obtain survival data spanning from 100% to 0%. LTe_50_ from workers as well as their 95% CI were extracted from fitted sigmoidal curves on survival data. Overall, cold-acclimation exerted a significant effect on LTe_50_ values across all species. The onset of colony-level dormancy also exerted a significant effect on LTe_50_ values, although to a lesser degree than acclimation, except for *L. flavus*. No differences in LTe_50_ values were observed between acclimated and acclimated + dormancy workers, except for *F. fusca*. Statistical significance between treatments within species is indicated at p < 0.05* (non-overlapping 95% CI).

### Colony-level dormancy aligns with suppressed metabolic rates in most, but not all, species

To confirm that the onset of colony-level dormancy coincided with suppressed metabolism indicative of diapause in individuals, we compared metabolic rates between workers from control and dormant conditions at 18°C (Fig. S2). The two-way ANOVA testing for the impact of species and dormancy on SMR values (Table S2) returned both factors and their interaction term as significant (p<0.001), indicating that across the global dataset and all species tested, dormancy had a significant effect on SMR values. A Tukey post-hoc test to assess pairwise differences specifically between dormant and active conditions within each species indicated, however, a statistically significant variation between SMR for three species only (*Lasius flavus*, *Formica fusca*, and *Myrmica rubra*), while no statistical differences in SMR were observed between control and dormancy for *Lasius niger* and *Leptothorax acervorum* (Fig. S2).

### Cold acclimation, but not colony-level dormancy, drives metabolomic shifts in workers

A heatmap representation of metabolites’ fold-change (FC) value between species and conditions (Fig. 4) supported that colony-level dormancy exerted only limited effect on workers’ metabolic profiles (Fig. 4A, D). In contrast, cold acclimation exerted a stronger effect on workers’ metabolomic profiles (Fig. 4B, C) with notable changes in the prevalence of some specific metabolites (see below). To analyze this metabolomic response more holistically, we performed a Principal Component Analysis (PCA). This PCA examined the patterns of relative metabolite abundance changes (area under the chromatographic peak) across our four experimental conditions and all species tested within a single analysis, incorporating CT_min_ and LTe_50_ values to assess potential alignment between increased cold hardiness and shift in metabolomic profiles of workers (Fig. 5A). The PCA analysis revealed that the first two principal components (PC) explained 24.5% and 15.5% of the total variance between conditions, respectively (Fig. 5A). CT_min_ and LTe_50_ strongly aligned with PC2, to which the four most contributing metabolites were glycerophosphocholine, glycerophosphoethanolamine, trehalose and inositol (Fig. 5B, Fig. S3). Average FC for these metabolites across cold-acclimated conditions were 8.73, 4.91, 2.28 and 1.54 as compared to non-acclimated conditions, respectively (Fig. 4). Other metabolites contributing to PC2 by decreasing relative contribution were D and L glutamic acid, Aspartic acid, Fumaric acid, Proline, Ribitol (Adonitol), Malic acid, Tryptophan, Lysine, Tyrosine. Together, these 13 metabolites accounted for 50% of the dataset contribution to PC2 (Fig. 5B, Fig. S3). A focused graphical representation of metabolite changes due to cold acclimation in species, mapped on schematic pathways of intermediary metabolism, is available in Fig. S4. The 13 metabolites contributing to up to 50% of PC2 are mapped in red (increased relative abundance) and blue (decreased relative abundance), respectively.

**Figure 4.**
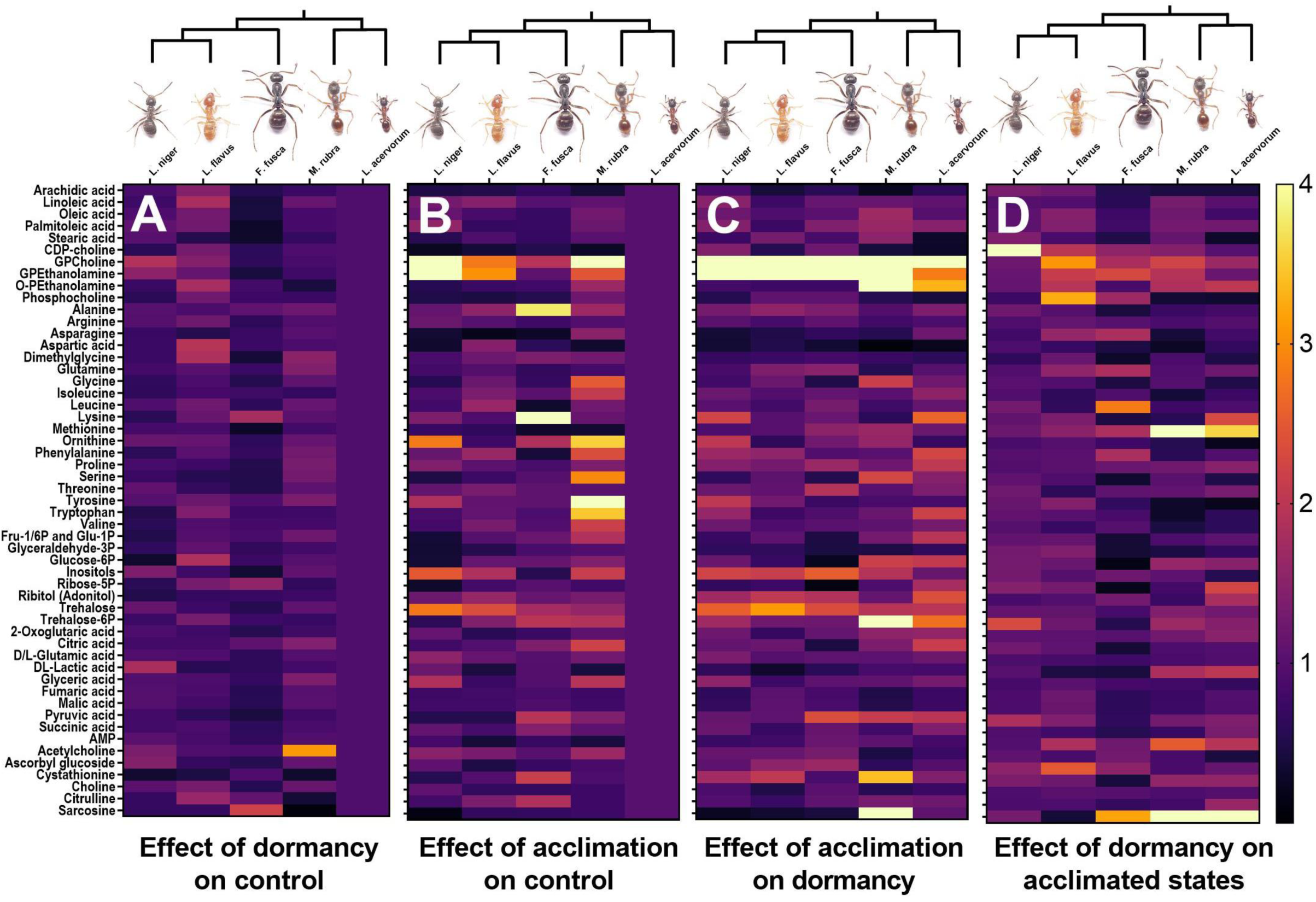
Heatmap visualization of metabolomic reorganization in workers. Data is structured to display the fold-change (FC) in metabolite area under the curve due to (A) the onset of colony-level dormancy on control workers, (B) the effect of cold acclimation on control workers, (C) the effect of cold acclimation on colony-level dormancy, and (D) the comparison between control-acclimated and diapausing-acclimated workers, singling out the effect of dormancy on acclimated states. Metabolomic data were deficient for *L. acervorum* in the control condition, resulting in metabolites’ FCs being fixed as 1 for this species as displayed in A and B. Overall, the onset of colony-level dormancy exerted a limited effect on metabolomic reorganization in species (A, D). In contrast, cold acclimation (B, C) was associated with more significant and consistent changes across profiles, with glycerophosphocholine, glycerophosphoethanolamine, and trehalose appearing to visually display the highest FC compared to control/dormancy conditions (also see PCA results in Fig. 4). A simplified comprehensive map of intermediate metabolism is provided in Fig. S5

**Figure 5.**
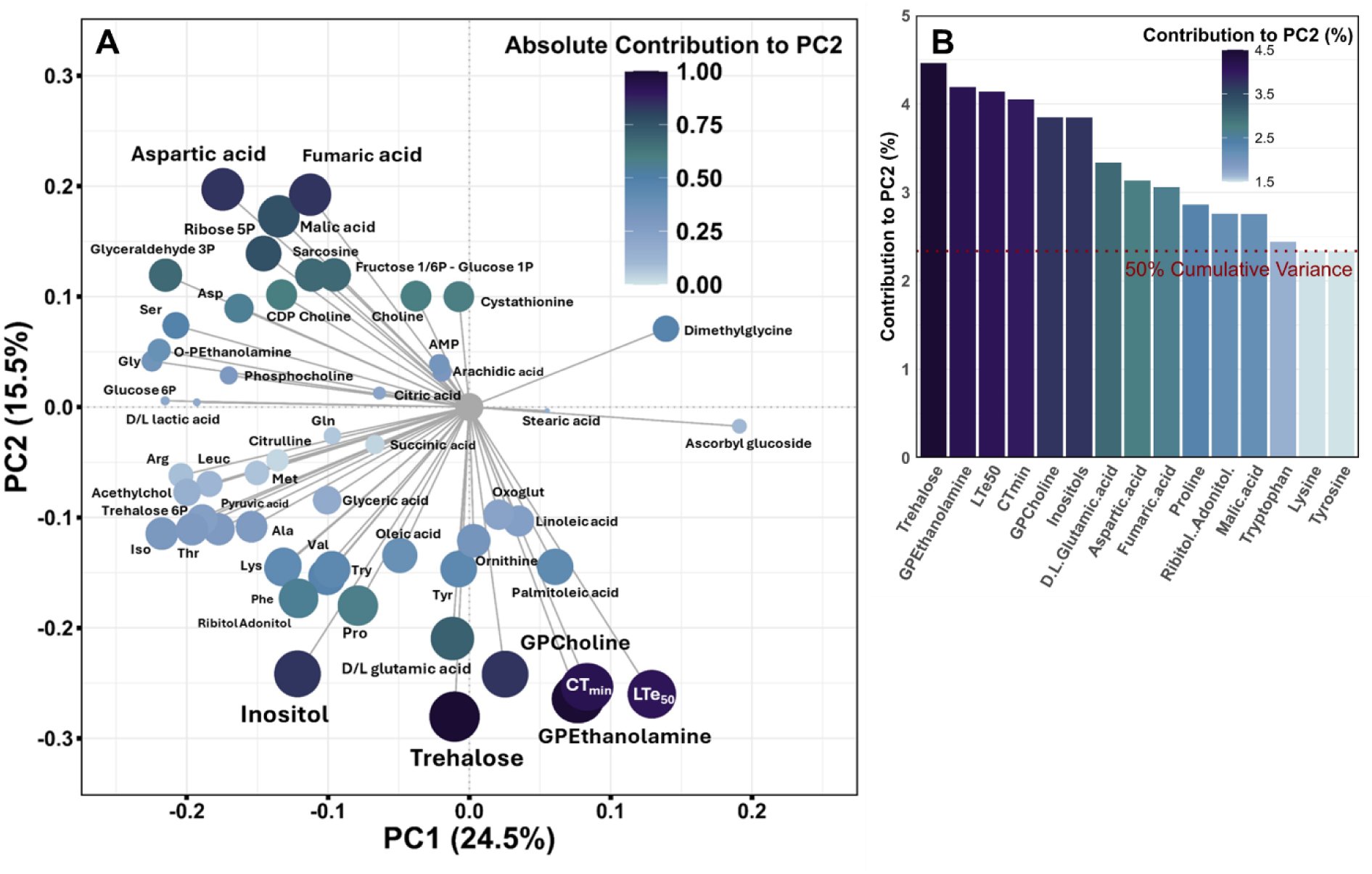
**A.** Principal Component Analysis (PCA) of metabolomic data, showing that variations of CT_min_ and LTe_50_ across treatments strongly aligned with PC2, to which the 4 most contributing metabolites were trehalose, glycerophosphoethanolamine, glycerophosphocholine, and inositol. **B.** Scree plot depicting the relative contribution of each metabolite and cold-hardiness variable to PC2 up to 50% of cumulative variance (a complete plot detailing the relative contribution of our 53 metabolites to PC2 is presented in Fig. S4). A simplified overview of the schematic pathways of intermediary metabolism impacted by cold acclimation is available in Fig. S5.

## Discussion

This work uses the endogenous developmental cycle of temperate ant colonies as a framework for understanding the relative contributions of colony-level dormancy and cold acclimation to cold tolerance and seasonal adaptation in ants (Fig. 1A, B). In endogenous-heterodynamic ants that display obligate colony-level dormancy, most individuals can experience multiple obligate diapause/colony-level dormancies over their lifespans^22–24^. Many species (including *Lasius*, *Myrmica*, and *Leptothorax* in this work) also exhibit brood development that can extend across more than one developmental season, leading to the presence of diapausing larvae within dormant colonies^22,24,37^. In contrast, other species (such as those from the genus *Formica,* e.g., *Formica fusca* in this work), complete their brood-rearing cycle within a brief annual period^22,24^, and colonies enter dormancy without brood^22,25,27^. Within our experimental system, we therefore used the combined cessation of worker foraging activity, the absence of queen egg laying, and the presence of arrested-development larvae (or complete brood absence in *F. fusca*) to define the onset of “colony-level dormancy”. Suppressed SMRs are also usually indicative of diapause in insects^7^. Across our dataset, we found support for a trend indicative of preprogrammed diapause associated with reduced SMRs of individuals and colony-level dormancy (Table S2), though this was not statistically significant for all species (Fig. S2). Together, our design thus offered a strong conceptual approach teasing apart the relative contributions of colony-level dormancy and acclimation on species’ cold tolerance.

Insects’ cold tolerance is typically quantified from one or more complementary traits^31,38^. Here, we integrated the use of CT_min_ and LTe_50_ values to characterize the cold tolerance of workers. CT_min_ is defined as the lower temperature at which individuals lose neuromuscular function and enter chill coma^39^. In insects, the onset of chill coma appears primarily driven by spreading depolarization of the central nervous system due to the loss of ion homeostasis within the CNS^40–42^. By contrast, LTe₅₀ is a measurement of cold mortality caused by homeostatic failure of the whole body; captured here by the temperature at which 50% of individuals die after a 24-h period of cold exposure^31^. Temperate ants are categorized as freeze sensitive, chill-tolerant insects: they are able to tolerate relatively low subzero temperatures as long as internal ice formation is prevented, but will ultimately experience mortality due to the direct and indirect effects of cold on homeostatic function above the freezing points of their tissues ^37,43–45^. In our work, CT_min_ thus captured the lower functional temperature for neuromuscular activity (reversible, non-lethal chill coma), while LTe₅₀ captured mortality limits associated with whole body chill-injury.

First, focusing on CT_min_, our results highlight a novel pattern for insect literature. The onset of colony-level dormancy alone was enough to significantly lower CT_min_ values (Fig. 2, Table 1). Moreover, the onset of colony-level dormancy displayed a strong interactive effect with acclimation to impact CT_min_ values further. This suggests two things. First, the onset of colony-level dormancy in endogenous-heterodynamic ants (to a degree associated with diapause, Fig. S2, Table S2) exerted significant phenotypic modification in workers modulating temperature activity thresholds. Second, that the onset of this “winter phenotype” potentiates workers to be more responsive to subsequent acclimation (Fig. 2, Table 1). CT_min_ values in insects and ants are documented both highly plastic and locally adaptive across populations and species^30,46–49^. Colonies studied here were collected in Aarhus, Denmark (56.16°N, 10.20°E), a region with a temperate oceanic climate and long, mild, winters averaging minimum temperature of ca. –0.56°C in January (1991-2023)^50^. CT_min_ values of dormant and cold-acclimated individuals (ranging from –3.2°C to –4.3°C; Fig. 2) closely matched the local winter air minima by a few degrees margin, resulting in a relatively low thermal safety margin when considering this trait. This suggests that workers may retain capacity for activity throughout much of the cold season, especially when relying on behavioral avoidance strategies via their thermally buffered subterranean nests.

Contrasting to CT_min_, the effect of colony-level dormancy on LTe₅₀ values was less consistent (Fig. 3, Table 1). This indicates that endogenous colony-level cycles had a less pronounced (but still detectable) effect on acute cold tolerance in most species. As expected, cold acclimation, on the other hand, consistently increased workers’ acute cold tolerance (Fig. 3, Table 1). LTe₅₀ values in acclimated workers ranged from –8.5°C to –13.5°C, indicative of significant capacity for surviving temperatures well below local winter minima in Denmark^50^. Our results thus illustrates a dual thermal safety margin strategy depending on the cold-tolerance trait considered: a narrow margin for activity (CT_min_), as previously documented in temperate insects^51^, but a much broader margin for lethal limits (LTe₅₀), maximizing survival capabilities. These findings underpin the importance of assessing multiple physiological traits when evaluating the ecological relevance of species’ cold tolerance, as different measures capture different aspects of thermal adaptation in ectotherms^31^.

Furthermore, mortality from acute cold-stress exposure (LTe₅₀) in chill-tolerant insects like ants is partly related to their ability to supercool their body fluids through the accumulation of cryoprotective metabolites^52^. To explore whether such biochemical adjustments could underpin shifts in LTe_50_ in heterodynamic ants, we conducted a targeted metabolomic analysis in workers following dormancy/acclimation. In this context, cold acclimation triggered a modest but consistent metabolomic shift in workers (Fig. 4, B, C), while colony-level dormancy had a more limited impact (Fig. 4, A, D). The metabolites most responsive to cold acclimation across our dataset were (i) two intermediates of phospholipid catabolism (glycerophosphoethanolamine and glycerophosphocholine) and (ii) two sugars (trehalose and inositol, Fig. 4,5, Fig. S3). However, fold-changes in these metabolites remained relatively modest comparative to other studies investigating metabolomics reorganization to cold in insects^53,54^. This could reflect the milder acclimation treatment applied to colonies (8°C for 10 days), which may not have been sufficient to trigger stronger physiological responses. A tentative interpretation could link the observed modulation of phospholipid catabolism to biological membrane remodeling. Membrane fluidity is highly temperature sensitive, and its disruptions can contribute to osmotic and ionic imbalance leading to cold-induced neuromuscular paralysis^42,55,56^. Second, polyol accumulation can act through multiple protective pathways. At high concentration, they can limit ice formation and lower the supercooling points of tissues, but can also be involved in stabilizing membranes and support osmotic balance^57–63^. Their modest upregulation in our work is likely indicative of osmotic and ionic homeostasis roles, rather than direct colligative depression of supercooling points, as previously suggested in *Drosophila*^32,64^. Polyol accumulation (e.g., glycerol) was previously reported in wintering cold-tolerant species of ants as well^37,43–45^. Further work would be needed to fully elucidate their role in the cold-tolerance strategies of temperate species.

In summary, our findings document the occurrence of endogenous cold hardening in seasonally adapted ants. These results provide the first evidence that thermal tolerance in social insects can be partly regulated by seasonal endogenous cycles, independently of environmental cues. They highlight the interplay between intrinsic seasonal programming and extrinsic thermal cues in shaping cold tolerance in temperate ants, offering insights into some of the key processes that have allowed this family of insects to thrive in highly seasonal climates.

## Supporting information

Electronic Supplementary Material

## Authors’ contributions

Q.W., VK and J.O. conceived and planned the study. Q.W. collected samples. Q.W. performed lab experiments. Q.W., and V. K analyzed the data. All authors contributed to drafting the article and approved the final published version.

## Acknowledgements

We thank L. Jørgensen for her help in taking care of captive ant colonies.

