## Supplementary material for "Endogenous circannual cycles drive seasonal cold hardening in temperate ants": Electronic Supplementary Material

Quentin Willot[1,2*], Vladimír Koštál[3], Johannes Overgaard[2]

[1]Research Unit of Environmental and Evolutionary Biology, Institute of Life, Earth & Environment, University of Namur, Namur, Belgium

[2]Section for Zoophysiology, Department of Biology, Aarhus University, 8000 Aarhus C, Denmark

[3]Institute of Entomology, Biology Centre, Czech Academy of Sciences, Branišovská, 1160/31, 37005 České Budějovice, Czech Republic

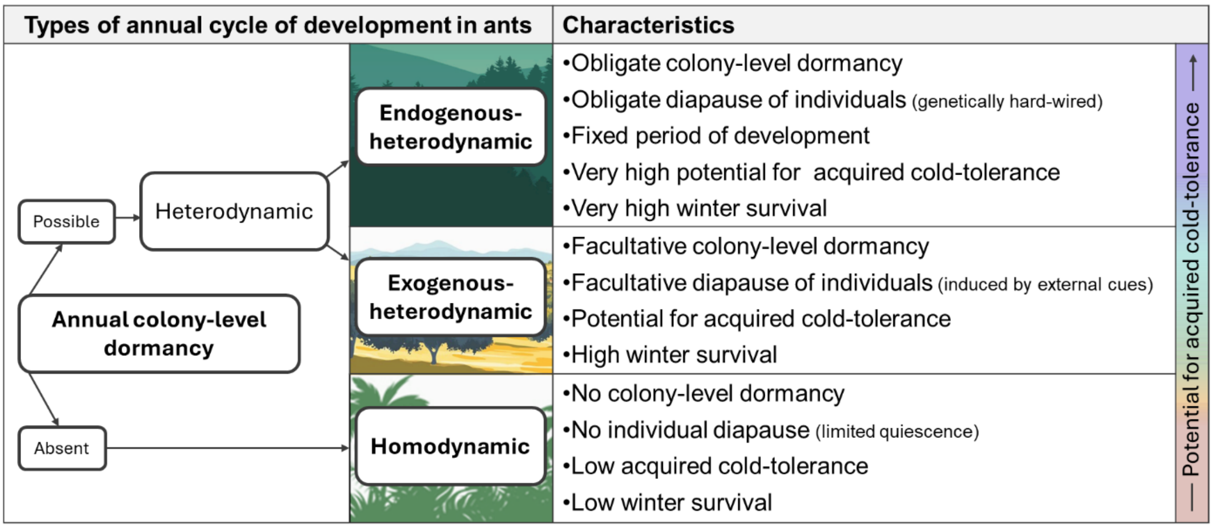


**Figure S1.** Simplified classification of seasonal life cycles in ants. Homodynamic ants, typically tropical, cannot halt colony growth in response to environmental cues and show limited acquired cold tolerance. In contrast, heterodynamic ants can experience periods of colony dormancy and suspended development. In exogenous-heterodynamic species, colony-level dormancy is triggered by environmental cues (facultative diapause), a strategy often observed in species adapted to subtropical/mediterranean climates and milder seasonal conditions. In endogenous-heterodynamic ants, dormancy and diapause are genetically programmed (obligate), occurring after a set period controlled by Kipyatkov's "sand glass device." This strategy is common in temperate and boreal species adapted to highly seasonal climates.

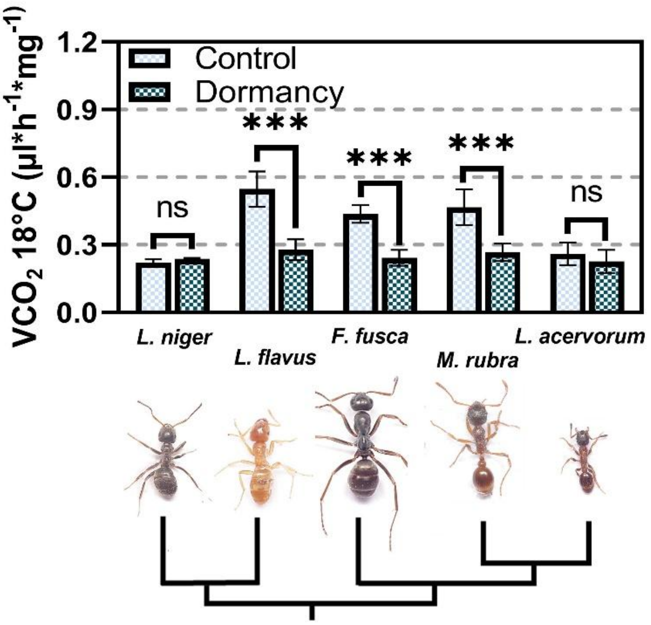


**Figure S2.** Standard Metabolic Rates (SMR) of workers at 18°C between control (workers from developing conditions) and dormant (workers from colonies having entered colony-level dormancy) conditions. SMR values were acquired through stop-flow respirometry on pooled individuals (15-20 ants), with 4 to 6 replicates per condition. No changes in experimental keeping conditions (12:12 light/dark cycles and 26°C) nor cold acclimation had taken place between control and dormant conditions. The two-way ANOVA testing for the impact of both species and dormancy on SMR values returned both factors and their interaction term as significant ($p<0.001$, Table S2), indicating that across the global dataset and all species tested, dormancy significantly impacted SMR values. However, the Tukey post-hoc test to assess pairwise differences specifically between dormant and active conditions within each species indicated statistical level of variation between SMR for three species only ($p<0.05$*, $p<0.01$**, $p<0.001$***).

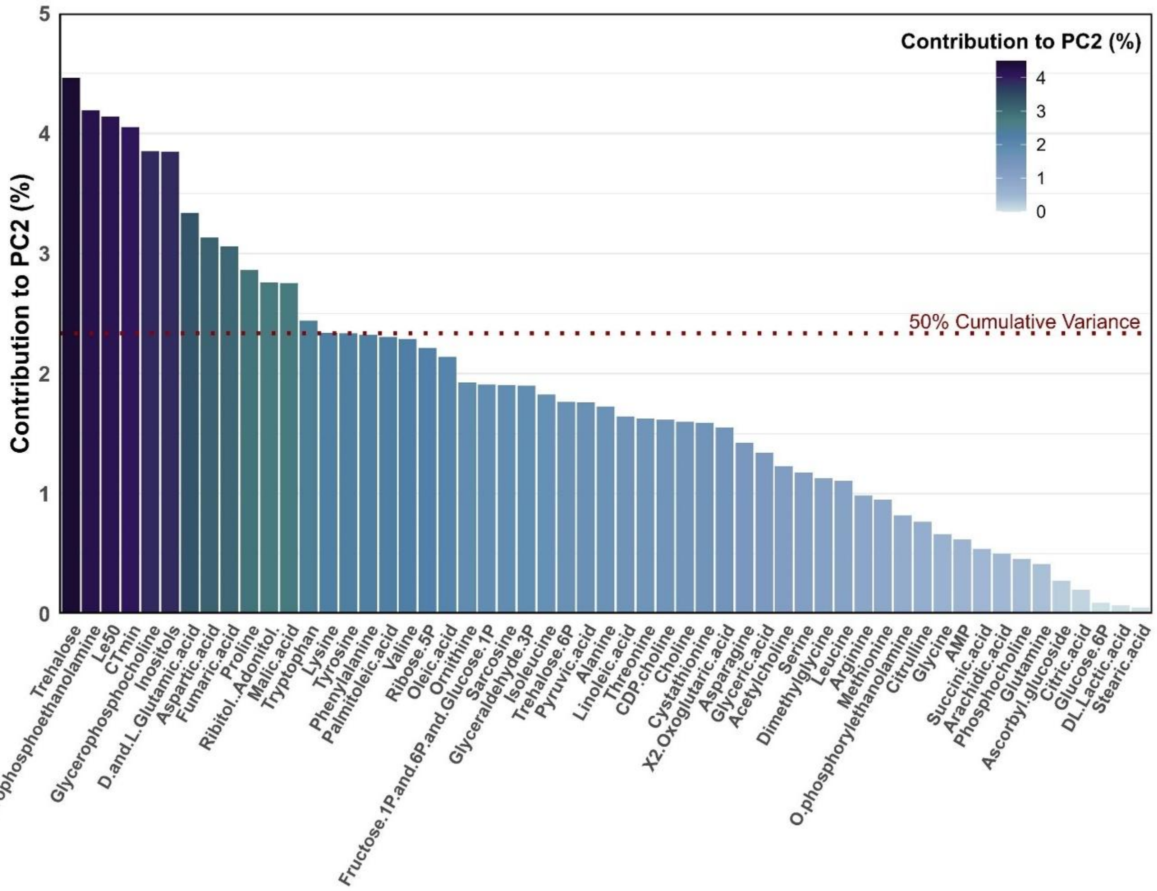


**Figure S3.** Scree plot depicting the relative contribution of each of the 53 metabolite and tolerance parameters variables included in the dataset to PC2. Fifty percent of the cumulative variance of PC2 was shared between 12 metabolites, of which trehalose, glycerophosphoethanolamine, glycerophosphocholine, and inositol were the top contributors.

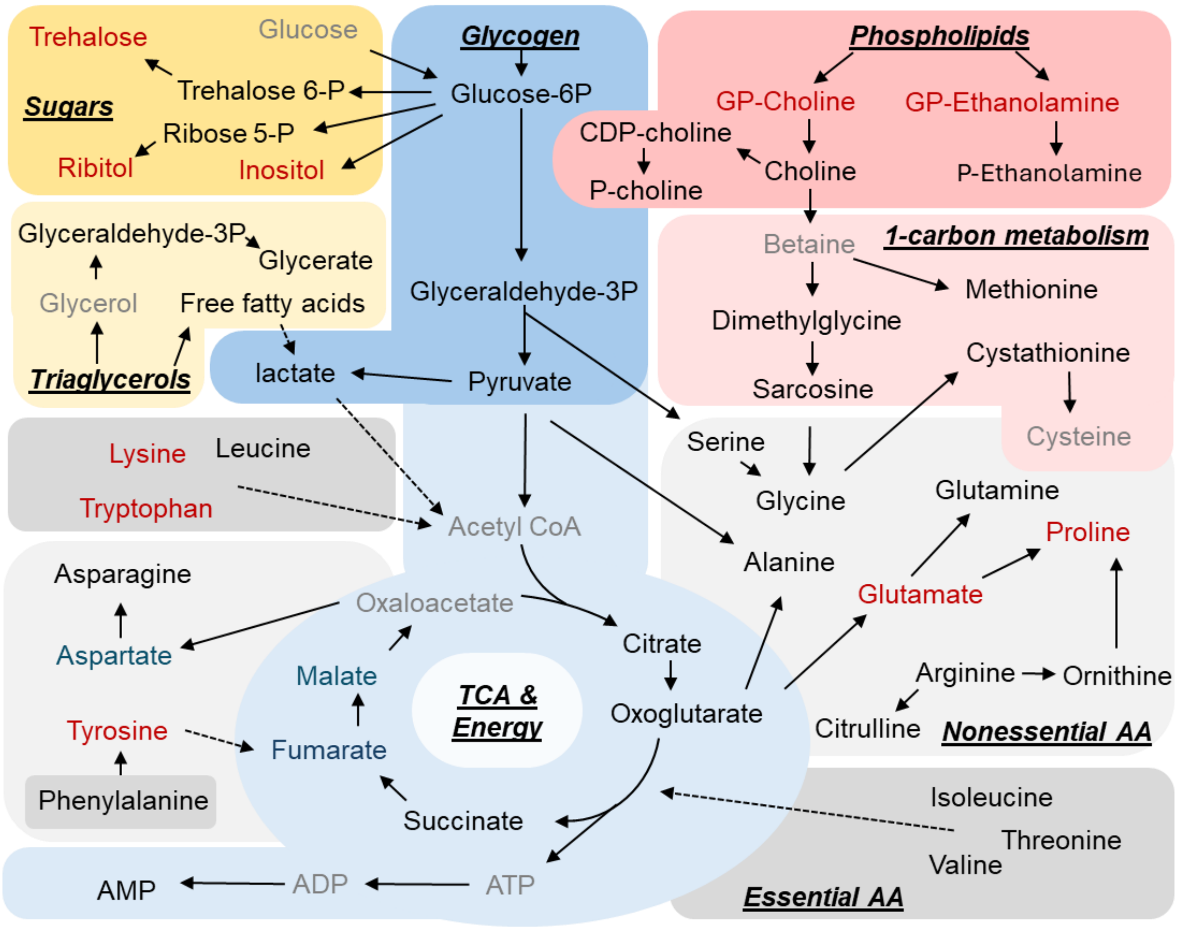


**Figure S4.** Simplified graphical representation focused on metabolite changes impacted by cold acclimation in species, mapped on schematic pathways of intermediary metabolism (pathways are indicated in italic, bold and underscored font separated within different color boxes). The 13 metabolites contributing up to 50% of PC2 (Fig. 4B, Fig. S5) are mapped in red (increased relative abundance) and blue (decreased relative abundance), respectively. Metabolites that were not detected or retained for analysis after QC (see methods) are greyed out. Other metabolites are shown in black. Broken arrows imply several biochemical steps between two connected metabolites.

| $CT_{min}$ model | $R^2$ | Delta (AIC) | AIC weight |
| --- | --- | --- | --- |
| $CT_{min}$ ~ 1 (null model) | 0 | 37.84 | $3.10*10^{-9}$ |
| $CT_{min}$ ~ acclimation + (1 \| sp) | 0.64 | 25.24 | $1.68*10^{-6}$ |
| $CT_{min}$ ~ dormancy + (1 \| sp) | 0.25 | 32.80 | $3.86*10^{-8}$ |
| $CT_{min}$ ~ acclimation + dormancy + (1 \| sp) | 0.94 | 0.10 | 0.49 |
| $CT_{min}$ ~ acclimation * dormancy + (1 \| sp) | 0.96 | 0.00 | 0.51 |
| $LTe_{50}$ model | $R^2$ | Delta (AIC) | AIC weight |
| $Lte_{50}$ ~ 1 (null model) | 0 | 10.31 | $4.34*10^{-3}$ |
| $Lte_{50}$ ~ acclimation + (1 \| sp) | 0.64 | 3.00 | 0.17 |
| $Lte_{50}$ ~ dormancy + (1 \| sp) | 0.17 | 9.09 | $7.94*10^{-3}$ |
| $Lte_{50}$ ~ acclimation + dormancy + (1 \| sp) | 0.71 | 0.00 | 0.74 |
| $Lte_{50}$ ~ acclimation * dormancy + (1 \| sp) | 0.70 | 4.71 | 0.07 |

**Table S1.** Coefficient of Determination ($R^2$) and Akaike information criterion (AIC) weights for each linear mixed-effects regression models tested to predict the impact of acclimation and colony-level dormancy on $CT_{min}$ and $LTe_{50}$ values. Values for models supporting the best prediction (Delta AIC = 0.00) are highlighted in blue. This indicates that the best model predicting $CT_{min}$ values integrated both acclimation and dormancy as well as their interaction as fixed effects. The best model predicting $LTe_{50}$ integrated both acclimation and dormancy as fixed effects (although dormancy returned as non-significant, see Table 1) but did not include their interaction.

| Two way ANOVA model | Factor | $p$ |
|---|---|---|
| SMR ~ dormancy * species | Dormancy | $9.10*10^{-13***}$ |
| | Species | $2.00*10^{-11***}$ |
| | Dormancy * Species | $9.17*10^{-10***}$ |

**Table S2.** Two-way ANOVA testing the effects of dormancy, species, and their interaction on the Standard Metabolic Rate (SMR) of workers. All factors were highly significant overall, indicating that both dormancy and species independently affected SMR. Additionally, the significant interaction term suggests that the effect of dormancy on SMR varied across species (see Fig. S2). A Tukey post hoc test assessing pairwise differences between dormant and active states within each species revealed significant differences in SMR due to dormancy for three species only (Fig. S2, $p < 0.05$*, $p < 0.01$**, $p < 0.001$***).
